# NERO: A Biomedical Named-entity (Recognition) Ontology with a Large, Annotated Corpus Reveals Meaningful Associations Through Text Embedding

**DOI:** 10.1101/2020.11.05.368969

**Authors:** Kanix Wang, Robert Stevens, Halima Alachram, Yu Li, Larisa Soldatova, Ross King, Sophia Ananiadou, Maolin Li, Fenia Christopoulou, Jose Luis Ambite, Sahil Garg, Ulf Hermjakob, Daniel Marcu, Emily Sheng, Tim Beißbarth, Edgar Wingender, Aram Galstyan, Xin Gao, Brendan Chambers, Bohdan B. Khomtchouk, James A. Evans, Andrey Rzhetsky

## Abstract

Machine reading is essential for unlocking valuable knowledge contained in the millions of existing biomedical documents. Over the last two decades ^1,2^, the most dramatic advances in machine-reading have followed in the wake of critical corpus development^3^. Large, well-annotated corpora have been associated with punctuated advances in machine reading methodology and automated knowledge extraction systems in the same way that ImageNet ^4^ was fundamental for developing machine vision techniques. This study contributes six components to an advanced, named-entity analysis tool for biomedicine: (a) a new, Named-Entity Recognition Ontology (NERO) developed specifically for describing entities in biomedical texts, which accounts for diverse levels of ambiguity, bridging the scientific sublanguages of molecular biology, genetics, biochemistry, and medicine; (b) detailed guidelines for human experts annotating hundreds of named-entity classes; (c) pictographs for all named entities, to simplify the burden of annotation for curators; (d) an original, annotated corpus comprising 35,865 sentences, which encapsulate 190,679 named entities and 43,438 events connecting two or more entities; (e) validated, off-the-shelf, named-entity recognition automated extraction, and; (f) embedding models that demonstrate the promise of biomedical associations embedded within this corpus.

---

Even the relatively specialized subfields of present-day biology and medicine are facing a deluge of accumulating research articles, patents, and white papers. It is increasingly difficult to stay up-to-date with contemporary biomedicine without the use of sophisticated machine reading (MR) tools. MR tool development, in turn, has been limited by the availability of biomedical corpora carefully annotated by experts. This is especially true with respect to information extraction, such as named entity recognition and relation or event extraction. Although several corpora have been developed for specialized biomedical subdomains, the need for a corpus that can bridge biological, general scientific, environmental, and clinical scientific sub-languages is greater than ever before.

Unfortunately, the annotation of natural science texts is more challenging than in other domains. Biomedical language is replete with ambiguity distinct from that observed in news articles or informal text online. When a word or phrase’s semantic meaning is clearly separated (*the east bank of the Danube* versus *Deutsche Bank*), we can implement automated sense disambiguation using machine learning tools. In biomedical texts, however, alternative meanings are not always clearly separated. The problem is not that a phrase can refer to several distinct, real-world entities in different contexts, but that the scientists writing articles typically do not separate competing, close meanings. For example, in some biomedical contexts, a named entity may refer to a *gene* or a *protein* with nearly equal probability; for example, “a mutant hemoglobin *α_2_*” can refer to either a gene or a protein. If the author meant *gene-or-protein A*, and we force an annotator to choose either interpretation *gene A* or *protein A*, the resulting annotation is of limited utility because the choice between *gene* and *protein* is random if the meanings are equally likely based on context. Ideally, a specialized ontology of text entities would allow an annotator to choose the proper level of annotation granularity (*gene-or-protein*, in this example), minimizing the need for forced, random decisions. To the best of our knowledge, there is no biomedical ontology that meets the requirements for capturing semantic ambiguity. We aimed to fill this gap by developing a specialized, variable-level meaning resolution ontology, a carefully curated corpus, along with corpus annotation tools, and a collection of text embedding analyses to evaluate our annotated corpus.

Our new ontology, called NERO, short for Named-entity Recognition Ontology, attempts to minimize unwarranted, arbitrary annotative semantic label assignments in text entities, see Figure 1. NERO captures named entities, starting with most broad and vague concepts close to the taxonomy’s root, finishing with the most narrow and concrete concepts at the taxonomy’s leaves. Hence, *DomainEntity–and* all ambiguous semantic classes–correspond to NERO’s taxonomy root. The basic division thereafter is between *TextEntity* and *AbstractEntity*, where *TextEntity* further splits into *NamedEntity, NamedEntityGroup*, *Relationship*, and *Pronoun*. After *NamedEntity*, the hierarchy reflects that which is written in biological entity descriptions, rather than in those entities’ lexical representation. NERO defines ambiguous concepts, such as *GeneOrProtein*, which subsumes both *Gene* and *Protein* using the following axiom: *EquivalentTo: ‘Gene’ or ‘Protein.’* There are no biological entities that are either a gene or a protein, but there are lexical entities that can belong to either named entity class. NERO uses this pattern to express appropriate ambiguity regarding text entities, preserving uncertainty from the text. In this way, NERO classes represent textual instances and not the actual biological entities to which these instances refer. It is, therefore, straight-forward to link between the lexical and biological entity through a relationship such as *‘is about’*. So, the NERO class *Protein ‘is about’* some specific concept *‘protein’* in an ontology pointing to real biological entities, such as the Protein Ontology ^5^

**Figure 1.**
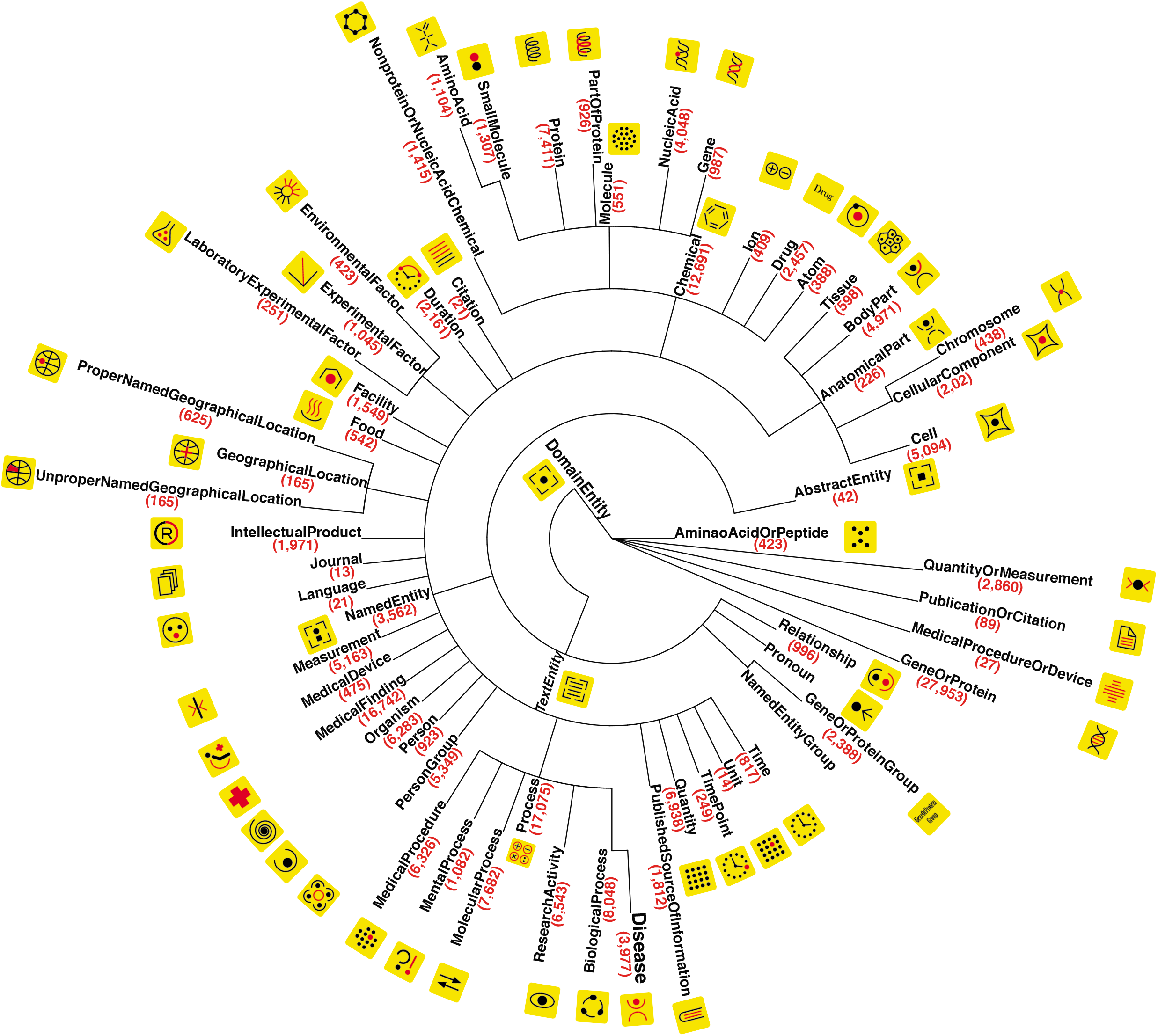
Named Entity Recognition Ontology (NERO). The Ontology is shown here as a multifurcating tree, with taxonomy nodes corresponding to ontology classes. Class name and class mentions count in the corpus are shown in parentheses next to each named entity class. Each taxonomy class is provided with a unique pictogram (black and red shapes on yellow background) intended to simplify expert manual annotation of the corpora. In total, we annotated 35,865 sentences. These sentences encapsulated 190,679 named entities and 43,438 events connecting two or more entities. In addition to the almost two dozen, more sparsely-used branches (such as *ExperimentalFactor* and *GeographicalLocation*) under the *NamedEntity* cluster, there are three heavily-represented branches in our corpus: *AnatomicalPart, Chemical*, and *Process*. Slightly more than half (51.6 percent) of all entities are from these three classes, with 26.6 percent of all entities originating from *Process* alone. We designed our ontology and its annotations to capture the named entities associated with research activities and facilities; these types of entities can be important for encoding methods used in scientific experiments or patient treatment. The semantic classes *ResearchActivity* and *MedicalProcedures* turn out to be the ninth and the tenth most frequent, respectively. Other top concepts related to research include *Measurement, IntellectualProducts, PublishedSourceOfInformation*, *Facility*, and *MentalProcess*.

Striving to make the ontology practically useful, we designed guidelines for annotators making decisions in annotating text entities, available in the *Supplementary Data*. Furthermore, by recruiting a team of postdoctoral-level experts, we annotated a large biomedical corpus to enable a broad range of natural language processing and biomedical machine learning tasks. Our annotations span 35,865 unique sentences, 8,650 of which were annotated by multiple annotators with remarkably high inter-annotator agreement (see Table 1). In our annotated corpus, we aimed to encompass all entity types that might occur in biomedical literature. In addition to named entities, our ontology captures *events* which represent relationships between biomedical concepts. The frequencies of all diverse entity types in our corpus are shown in Figure 2A; Figure 2B shows the frequencies of relations represented in the taxonomy. The most frequent entity type is *GeneOrProtein*, which accounts for 14.7 percent of all named entities in the corpus (see Figure 2A). The second most populous category is *Process*, with nine percent tagged. *Process* has six sub-concepts and almost half of *Process* instances (49.7 percent) are annotated as more specific sub-concepts; the *BiologicalProcess* and the *MolecularProcess* are the fifth and seventh most frequent entity types (see Figure 2). Entity type frequencies follow a heavy-tail distribution, with the least frequent types being *Journal, Unit*, and *Citation* (see Figure 2). In addition to 190,679 named entities, we annotated 43,438 action terms, events connecting two or more entities. The most annotated action term is *bind*, accounting for 28.4% of all actions, see *Supplementary Figure 1*. When we normalize the action terms and combine actions such as *bind, binds*, and *binding*, the normalized action *bind* accounts for 31.8% of all actions, as shown in *Supplementary Figure 1*. We deployed a package called NERO-nlp for researchers interested in diving deeper into our annotated corpus; the installation guides and scripts are available online at https://pypi.org/project/NERO-nlp and https://github.com/Bohdan-Khomtchouk/NERO-nlp respectively.

**Figure 2.**
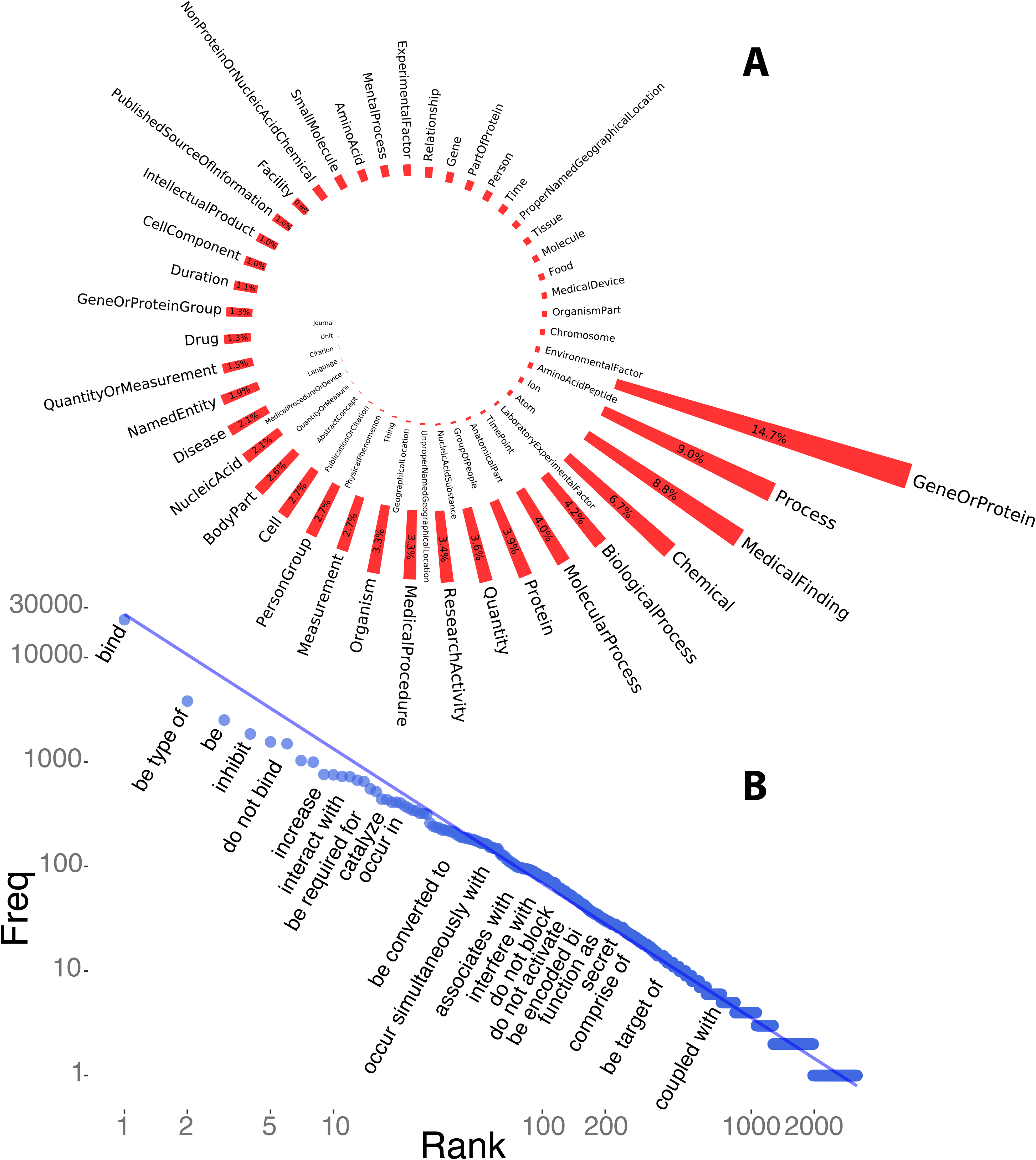
The relative abundance of annotated named entity classes in our corpus. As is typically the case with human languages, semantic classes are represented unevenly in free texts, following a heavy-tail (Zipf’s) distribution. (A) In biomedical corpora, unsurprisingly, named entities associated with *genes* and *proteins* are the most prevalent (15 percent), followed by *processes* (9 percent), *medical findings* (8.8 percent), and *chemicals* (6.7 percent). At the low-frequency end of the named entity spectrum, we find *journal names, units, citations*, and *languages*. (B) Events connecting two or more entities are also approximately Zipf-law distributed. Event frequencies are closely tracking corresponding named entity classes. For example, the most frequent event, *bind*, is associated with the most frequently named entity, *GeneOrProtein*. We tried fitting the rank-ordered frequency distribution of annotated named entities with a Discrete Generalized Beta Distribution (DGBD). The result showed a significant deviation from Zipf’s law The observed distribution’s tail was not heavy enough to match Zipf’s distribution, most likely due to the relatively small number of classes in our ontology. ^29^ In other words, we expect that frequencies of semantic classes in a very large corpus, annotated with classes from a hypothetical perfect named entity ontology, would follow a Zipfian (discrete Pareto) distribution of named entity classes. Our action annotations have moved beyond interactions between proteins and genes (*e.g., bind, inhibit*, *phosphorylate*, *encode*), into interactions involving genetic variants and environmental factors (*e.g*., *associated with, occur in presence of, trigger, lack*). Ambiguity levels varied broadly across the named entities captured in our corpus. For example, in the class *AnatomicalPart*, almost all (99.3 percent) are annotated at the most specific levels, with the majority of entities belonging to *BodyPart, CellularComponent*, and *Cell*. In contrast, the general (most vague) concept, *Chemical*, turns out to be the most annotated within its cluster, although more specific subclasses, such as *Protein, NucleicAcid*, and *Drug* are also well represented in the corpus. In the *Process* concept cluster, about a third of all concept instances are annotated at a more general *Process* level, and the rest of them are specific concepts, such as *MedicalProcedure*, *MolecularProcess*, *ResearchActivity*, and *BiologicalProcess*. In addition to these major clusters of concepts, several individual concepts are well represented in the corpus. For example, *MedicalFinding* represents 7.3 percent of all entities. Other well-represented concepts include *Duration, IntellectualProduct, Measurement*, *Organism*, *PersonGroup*, *PublishedSource OfInformation*, and *Quantity*. In total, about 70.4 percent of all entities are annotated at the most specific ontology level. There are five concepts in the NERO ontology that allow the semantic flexibility needed to avoid arbitrary concept assignment. Entities annotated as *AminaoAcidOrPeptide*, *QuantityOrMeasurement*, *PublicationOrCitation MedicalProcedureOrDevice*, and *GeneOrProtein* account for 17.8 percent of all entities, while less than a quarter (23 percent) of entities representing either genes or proteins are cleanly annotated with class *Gene* or class *Protein*. The remainder are annotated with class *GeneOrProtein*. In addition to the action *bind*, actions indicating entities’ attributes are the next most frequent. Other biological relationships are also well-represented in this annotation, such as *inhibit, activate, mediate, interact, contain*, and *regulate*. The top 30 action categories account for 64.4 percent of all actions annotated with the top ten action categories accounting for 52.2 percent. Interestingly, negations of actions were also quite abundant in our annotated corpus. For example, *do not bind* was the sixth most frequent normalized action. Other well-represented negations of actions include *do not affect* and *do not inhibit* (see *Supplementary Figure 1*).

**Table 1.**
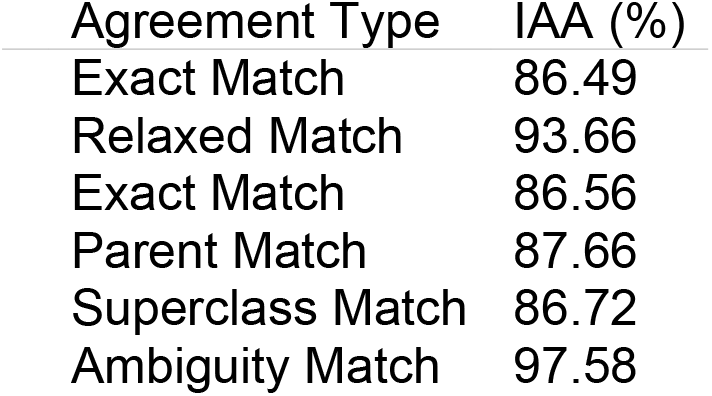
Inter-annotator Agreement Statistics.

| Agreement Type | IAA (%) |
| --- | --- |
| Exact Match | 86.49 |
| Relaxed Match | 93.66 |
| Exact Match | 86.56 |
| Parent Match | 87.66 |
| Superclass Match | 86.72 |
| Ambiguity Match | 97.58 |

Below, we present two practical applications of our ontology and text annotations: 1) Machine learning experiments, which automatically identify named entities, and; (2) Word embedding experiments, which leverage the automated discovery of semantic relationships among real-world concepts referenced by a text’s named entities.

## Machine learning experiments

Using NERsuite ^6^, we conducted a ten-fold cross-validation, dividing the corpus into training and test subsets. The classification results are presented in *Supplemental Table 1*. The overall automated named entity recognition performance is moderate, with 54.9% precision, 37.3% recall and a 43.4% *F*_1_ score. The best performance class, *GeneOrProtein*, had baseline results of 67.0% precision, 65.3% recall, and a 66.2% F_1_ score. In addition to the default baseline implementation of NERsuite, we added additional features in the training process to improve its performance ^7^ These are dictionary features derived from lookups in technical term dictionaries. The classifier with dictionary features manifests 54.7% precision, 37.9% recall and a 43.8% *F*_1_ score. We observed a scant 0.35% increase in *F*_1_ score from adding dictionary features. We then implemented an ensemble method called stacking, where we trained a higher-level model to learn how to best combine contributions from each base model. The base model in this case is the baseline model from NERsuite. Stacking yielded a 0.27% increase in *F*_1_ score compared to baseline results. While ensemble methods are commonly used to boost model accuracy by combining the predictions of multiple machine learning models, choices of second-level and base models can influence the amount of improvement in model accuracy. The overall performance statistics are shown in *Supplementary Table 2*. As our corpus is made public with this study’s publication, we hope that other researchers will use this training data to achieve core MR task performance that surpasses our initial experiments.

To examine how NERsuite performs in comparison to other popular open-source Named-Entity recognition tools, we trained a custom NER model on our annotated corpus using spaCy ^8^. We evaluated the trained model on the test subset, which consists of a random 10% sample from the corpus. Classification results are presented in *Table 3*. Overall automated named entity recognition performance is low, with 30.9% precision, 8.6% recall and a 13.4% *F*_1_ score. The best performance class, *GeneOrProtein*, had results of 45.1% precision, 36.4% recall, and a 40.3% *F*_1_ score. These statistics indicate a much poorer performance of spaCy compared to that of NERsuite.

**Table 3:** Experimental results using spaCy for NER evaluated on 10% of the corpus

|  | **p** | **r** | **f** |
| --- | --- | --- | --- |
| **Cell** | 16.88 | 5.10 | 7.83 |
| **CellComponent** | 35.71 | 4.44 | 7.91 |
| **GeneOrProtein** | 45.11 | 36.44 | 40.31 |
| **Organism** | 16.99 | 5.53 | 8.34 |
| **Disease** | 11.79 | 8.12 | 9.61 |
| **Drug** | 11.11 | 1.20 | 2.17 |
| **SmallMolecule** | 0.00 | 0.00 | 0.00 |
| **BiologicalProcess** | 8.02 | 1.88 | 3.05 |
| **MolecularProcess** | 12.67 | 2.45 | 4.10 |
| **Gene** | 0.00 | 0.00 | 0.00 |
| **Protein** | 0.00 | 0.00 | 0.00 |
| **BodyPart** | 15.17 | 5.54 | 8.12 |
| **AminoAcid** | 12.50 | 0.88 | 1.64 |

To help explain the huge difference in performance between NERsuite and spaCy, we considered the set of input features used by each tool for insight. NERsuite’s baseline implementation uses an extra set of input features including the lemma, POS-feature and chunk-feature, whereas our custom spaCy NER model only relies on character offsets and entity labels. There is potential for further customizing spaCy’s processing pipelines by adding more components such as tagger and parser ^8^, but no established approaches in this regard have been made available partly because spaCy’s model architecture is different from those of other popular NER tools. We also observed that some entities classes, such as Gene and Protein, have zero values for precisions, recalls and *F*_1_ scores, which likely translate to no correct classifications made for those entities. The zero values occur partly due to the relatively smaller number of tokens for those entity classes in the training set, and as a result, the trained NER model generalized poorly on the minority class entities in the test subset.

Due to spaCy’s computational demands, we did not conduct 10-fold cross-validation. NERsuite provides a well-integrated pipelined system where training a new model consists of a few lines of code. In addition, NERsuite has a demonstrated record ^5^ on two biomedical tasks, the BioCreative2 gene mention recognition task and the NLPBA 2004 named entity recognition task. Therefore, one could argue that it offers an advantage over spaCy for NLP tasks in specialized domains such as biomedicine.

We’ve also identified another package called scispaCy ^9^ that contains spaCy models for processing biomedical, scientific or clinical text. SciSpaCy acts as an extension to spaCy and provides a set of practical tools for text processing in the biomedical domain ^9^ In particular, scispaCy includes a set of spaCy NER models trained on popular biomedical corpora, which covers entity types such as chemicals, diseases, cell types, proteins and genes. As an extension to spaCy, it also has the flexibility for users to train a custom NER model from scratch or update the existing NER models with users’ own training data. Since our NER ontology adopts a more diverse and detailed annotation methodology for named entity types, it will be challenging to update scispaCy’s pretrained named entity recognizer with our annotated corpora.

## Word embedding experiments

Semantic associations, automatically extracted from text using neural network embedding operations, can function as a kind of “digital double” of real-world phenomena embedded in text, facilitating inferences that were previously imagined only possible from the original experimental data. For example, word embeddings built from chemical and material science texts predict much of the subsequent decades’ material discoveries^8^, just as the corpus of molecules can recover the periodic table^9^, and texts are able to recover the subtle, psychological and sociological biases of cultures that produced them ^10,11^. We used word embedding models to evaluate the biomedical veracity of NERO and its text annotation. Embedding models like Google’s `word2vec` ^12,13^ initially received substantial attention based on their capacity to solve analogy problems and automatically capture deep semantic relationships among concepts. Building on these capacities ^10,14,15^, we proposed a general method for constructing meaningful dimensions by taking the arithmetic mean of word vectors representing antonyms along a dimension and using them to diagnose their meanings. This approach has been widely validated ^15–19^, and we employed it here to construct and compare the meanings embedded in NERO and our annotated corpus with ground truth data about drugs and diseases. In order to evaluate word embeddings based on NERO, we identified two disease properties —(1) severity and (2) gender specificity—and likewise two therapeutic drug properties —(1) toxicity and (2) expense—not directly present in text, but highly relevant to diagnosis and treatment, and on which text-independent ground truth data exists.

We embedded named entities associated with diseases and drugs into a high-dimensional space in which every NERO term was assigned a 300-dimensional vector, (see Figure 3 for a three-dimensional projection of this embedding), along with a selection of diseases and medications used to treat them. We then compared drug and disease projections into the embedding dimensions for severity, gender, toxicity, and expense with ground truth about these qualities. We constructed the severe-mild axis with the following contrasting term pairs: (harmful, beneficial), (serious, benign), (life-altering, common), (disruptive, undisruptive), (dying, recovering), (dangerous, safe), (threatening, low-priority), (high mortality, harmless), (costly, cheap), (hospitalized, self-administered), (hospital, work), (debt, savings), (low quality of life, undisruptive), and (hazard, routine). Then we compared disease projection in this dimension with World Health Organization data on the burden of living with each of those diseases (DALYs ^20^) and found a correlation of 0.329 (*p*=0.0614, *n*=33). We then constructed a gender dimension with similarly contrasting pairs: (male, female), (prostate, ovary), (penile, uterine), (penis, uterus), (man, woman), (men, women), (masculine, feminine), (he, she), (him, her), (his, hers), (boy, girl), and (boys, girls). We compared the disease projection in this gender dimension with the prevalence of those diseases for men and women from a substantial sample of doctor-patient insurance records capturing approximately 47% of all of U.S. doctor-patient visits between 2003 and 2011 and found a correlation of 0.436 (*p*=1.46 × 10^−13^, *n*=261).

**Figure 3.**
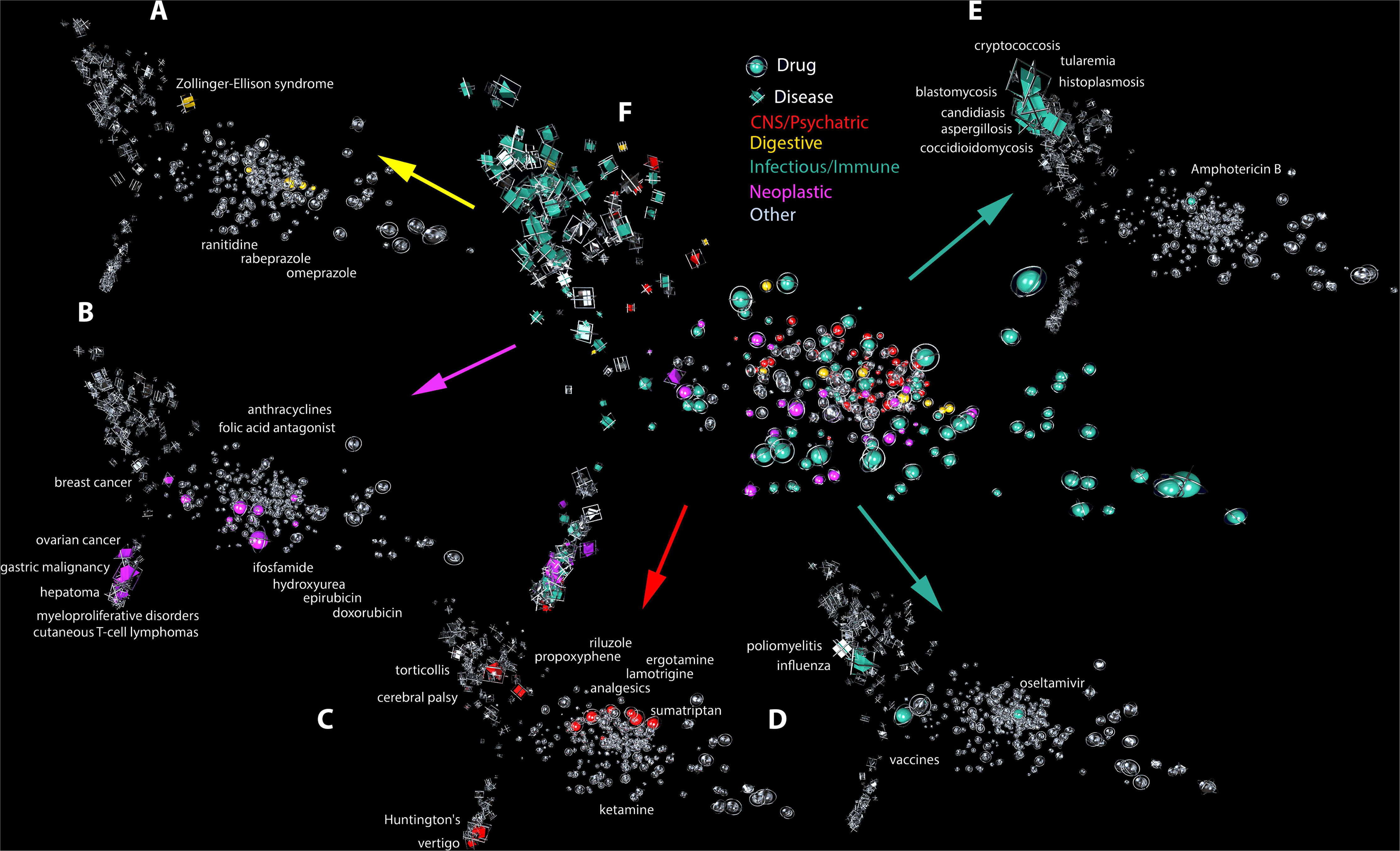
Properties of diseases and drugs visible in the first three principal components of our multi-dimensional text embedding. The figure shows a projection of text embedding into three-dimensional space, with named entities corresponding to diseases and drugs shown with prisms and spheres, respectively. The figure represents several projections of the same embedding, preserving spatial layout and projection, with distinct elements of the embedding indicated by shape color. The central image shows all disease systems and their corresponding medications together. More specifically, the additional projections show: (A) Zollinger-Ellison syndrome and associated medications; (B) cancers and associated therapies; (C) central nervous system diseases and corresponding medications, and; (D) and (E) Viral and bacterial infectious diseases, respectively, together with corresponding antiviral and antibiotic agents. Another view of the same dataset is presented in Figure 4.

Together, these patterns suggest that not only does NERO facilitate efficient and accurate concept-by-concept annotation, but that the distribution of biomedical properties underlying NERO-annotated texts have emergent validity and predict data patterns not explicitly present in biomedical articles. Following the same pattern, we projected medications onto a toxicity axis composed from: (harmful, beneficial), (toxic, nontoxic), and (noxious, benign) and an expense dimension anchored by: (expensive, inexpensive), (costly, cheap), (brand, generic), and (patented, off-patent). The correlation of drug projections onto the toxicity dimension correlates at 0.32 (*p*=1.1 × 10^−4^) with the median lethal dose, or dose required to kill 50% of subjects as documented in the LD50 database ^21^. Finally, the correlation of drug projections into an expense dimension and the price of each drug as listed in the IBM MarketScan database ^22^ was 0.42 (*p*=1.5 × 10^−15^) (see Figure 4). When a disease projects low in the *male – female* dimension, it is much more likely to afflict women than men, such as ornithosis and related infectious diseases. When a disease projects high in the *serious – benign* dimension like leprosy, it is likely to incur substantial suffering. When a medication projects high in the *toxic – nontoxic* dimension, such as Riluzole, a treatment for amyotrophic lateral sclerosis with potential severe side effects ranging from unusual bleeding to nausea and vomiting. Drug projections high in the *expensive – inexpensive* dimension suggest a stiff medical bill, as in the case of Simvastatin, which is used to reduce the risk of heart attack and stroke, and which, before it went off-patent, cost hundreds of dollars per bottle. The robust accuracy of these associations suggest that for qualities on which we do not have relevant or inexpensive data outside text, associations from text represent a significant signal for biomedical research and can be considered robust hypotheses meriting empirical study.

**Figure 4.**
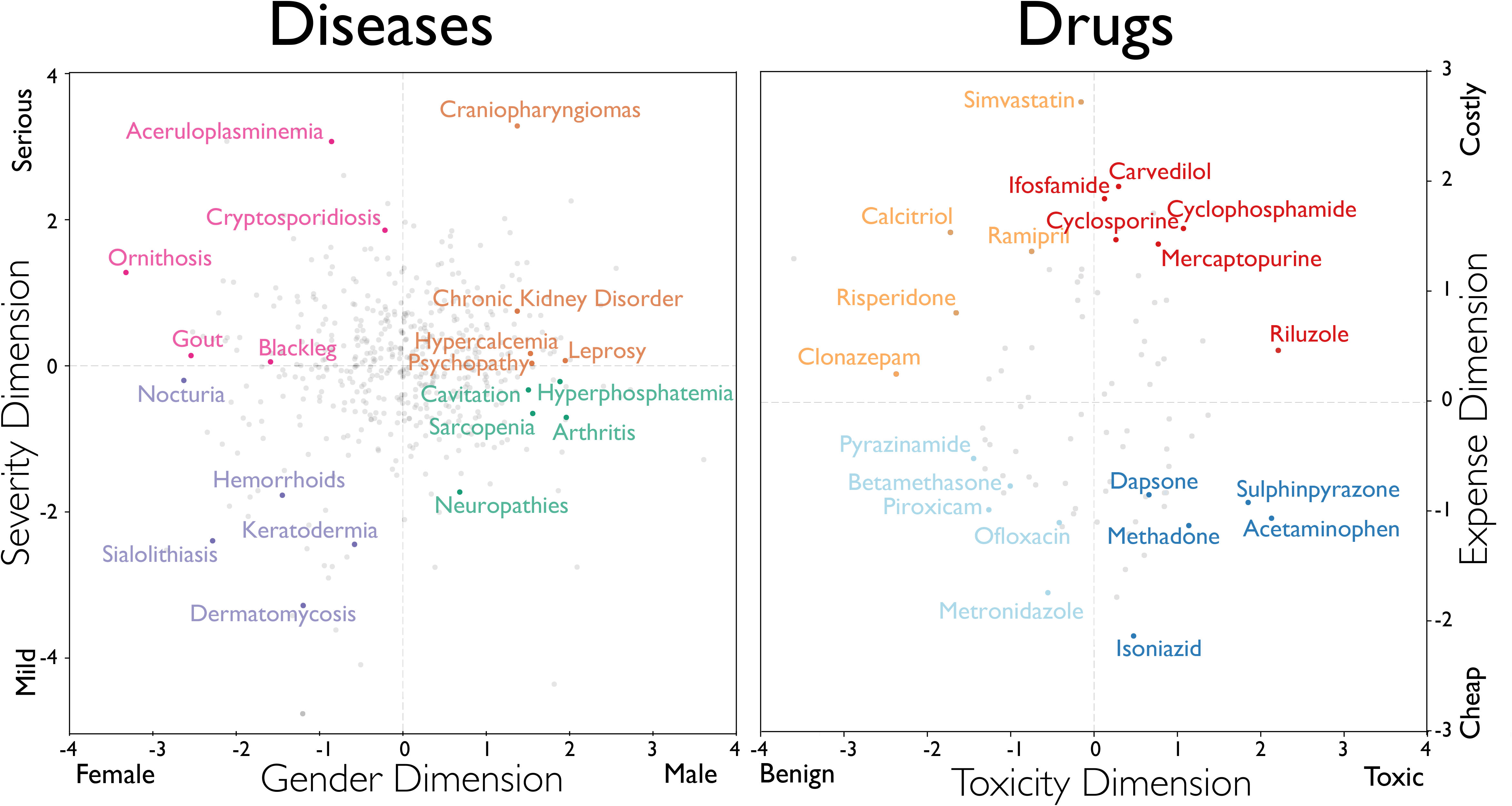
Two-dimensional projections of diseases and medications. (A) We projected diseases into two dimensions: female-male (X-axis) and severe-mild (Y-axis). We defined the “male-female” axis using the following pairs of terms: (‘male,’ ‘female’), (‘prostate,’ ‘ovary’), (‘penile,’ ‘uterine’), (‘penis,’ ‘uterus’), (‘man,’ ‘woman’), (‘men,’ ‘women’), (‘masculine,’ ‘feminine’), (‘he,’ ‘she’), (‘him,’ ‘her’), (‘his,’ ‘hers’), (‘boy,’ ‘girl’), and (‘boys,’ ‘girls’). We defined the severe-mild axis with the following term pairs: (‘harmful,’ ‘beneficial’), (‘serious,’ ‘benign’), (‘life-altering,’ ‘common’), (‘disruptive,’ ‘undisruptive’), (‘dying,’ ‘recovering’), (‘dangerous,’ ‘safe’), (‘threatening,’ ‘low-priority’), (‘high mortality,’ ‘harmless’), (‘costly,’ ‘cheap’), (‘hospitalized,’ ‘self-administered’), (‘hospital,’ ‘work’), (‘debt,’ ‘savings’), (‘low quality of life,’ ‘undisruptive’), and (‘hazard,’ ‘routine’). **(B)** We projected medications into “benign-toxic” (X-axis) and “cheap-costly” (Y-axis). For the “benign-toxic” axis, we used the following pairs of antonym words: (‘harmful,’ ‘beneficial’), (‘toxic,’ ‘nontoxic’), and (‘noxious,’ ‘benign’). We defined the “expensive-inexpensive” dimension using the following pairs of terms: (‘expensive,’ ‘inexpensive’), (‘costly,’ ‘cheap’), (‘brand,’ ‘generic’), and (‘patented,’ ‘off-patent’).

This study’s main limitation is that, even though our NERO ontology aimed to cover all entities contained in the biomedical research literature, we did not cover all levels of granularity in classifying entities. Moreover, while the major concepts are well-annotated, several concept types were not well-represented because of the heavy-tail distribution of ontological class frequencies. In addition, we note that satisfactory results of Named-entity Recognition (NER) rely heavily on a large quantity of hand-annotated data, which is often costly in terms of time and resources spent. Therefore, adoption of semi-supervised learning methods, which incorporates unlabeled data to improve learning accuracy, could reduce the need for manual annotation ^23^

While there is popular belief that pretraining on general-domain text can be helpful for developing domain-specific language models, a recent study has shown that for specialized domains, such as biomedicine, pretraining on in-domain text from scratch offers noticeable improvements in model accuracy compared to continual pretraining of general-domain language models ^24^ Therefore, we trained on our annotated corpus from scratch using in our machine learning experiments ^25^

The resources offered in our study can be applied to a wide range of scientific problems. First, the proposed NERO ontology can facilitate more robust and accurate large-scale text mining of biomedical literature. As discussed above, NERO is the first knowledge graph in this field, accounting for context-relevant levels of ambiguity. Graph neural networks ^26^ can leverage such prior knowledge from human experts for learning embedding of biomedical entities, which is likely to preserve both semantic meaning in the original literature and domain knowledge from human experts. Second, researchers can combine the curated corpus from this study with self-supervised learning ^27^ Such a learning scenario can utilize the unlabeled data in a supervised way by predicting part of the sentence using the rest of the sentence. The annotated corpus from this study can be used to fine-tune language models, orienting them for critical biomedical tasks.

## Supporting information

Ontology Corpus Annotation Guidelines

## Acknowledgments

We are grateful to E. Gannon and M. Rzhetsky, for comments on earlier versions of this manuscript. This work was funded by the DARPA Big Mechanism program under ARO contract W911NF1410333, by National Institutes of Health grants R01HL122712, 1P50MH094267, and U01HL108634-01, and by a gift from Liz and Kent Dauten. Additional support came from King Abdullah University of Science and Technology (KAUST), awards number FCS/1/4102-02-01, FCC/1/1976-26-01, REI/1/0018-01-01, and REI/1/4473-01-01.

