## Supplementary Figures for "NERO: A Biomedical Named-entity (Recognition) Ontology with a Large, Annotated Corpus Reveals Meaningful Associations Through Text Embedding"

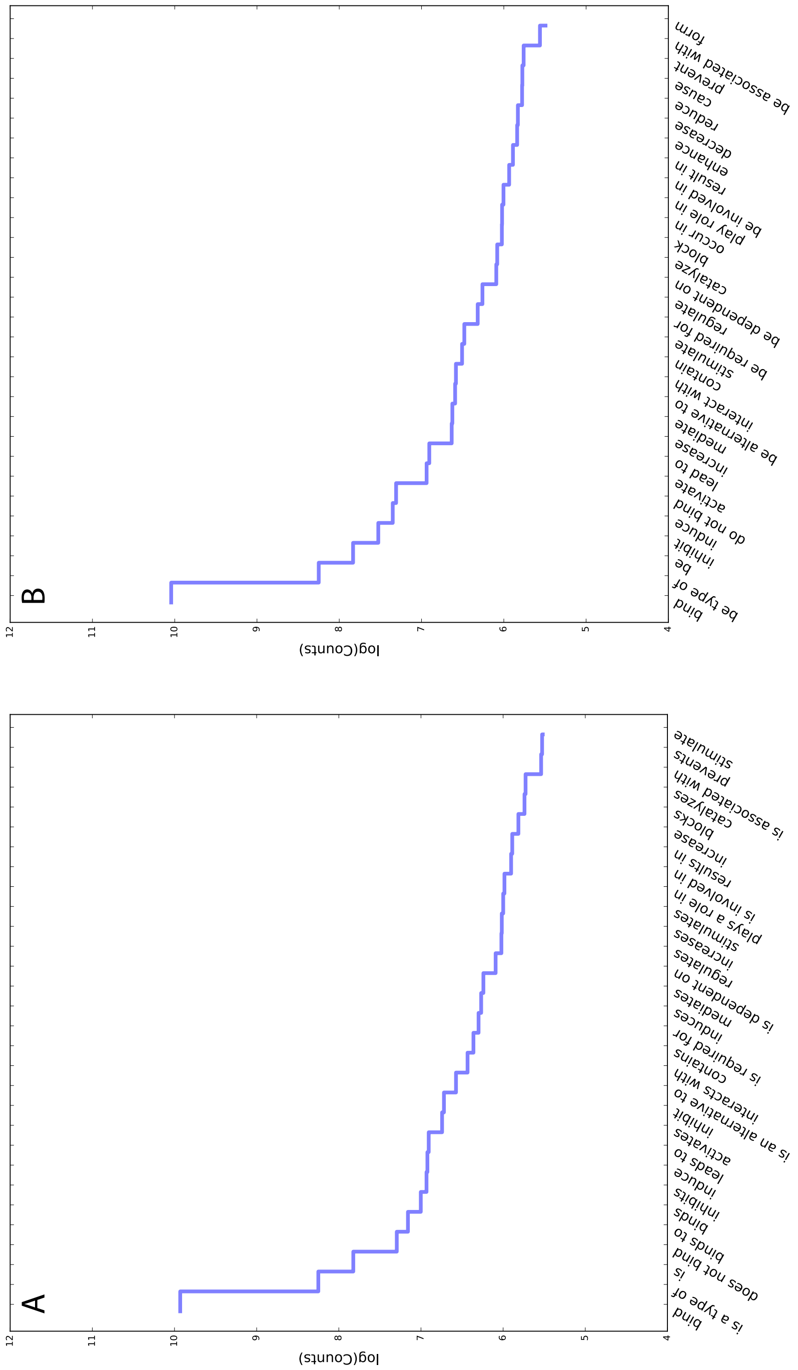


FIGURE 1. **Top 30 Actions** (A) Without normalization (B) With Normalization

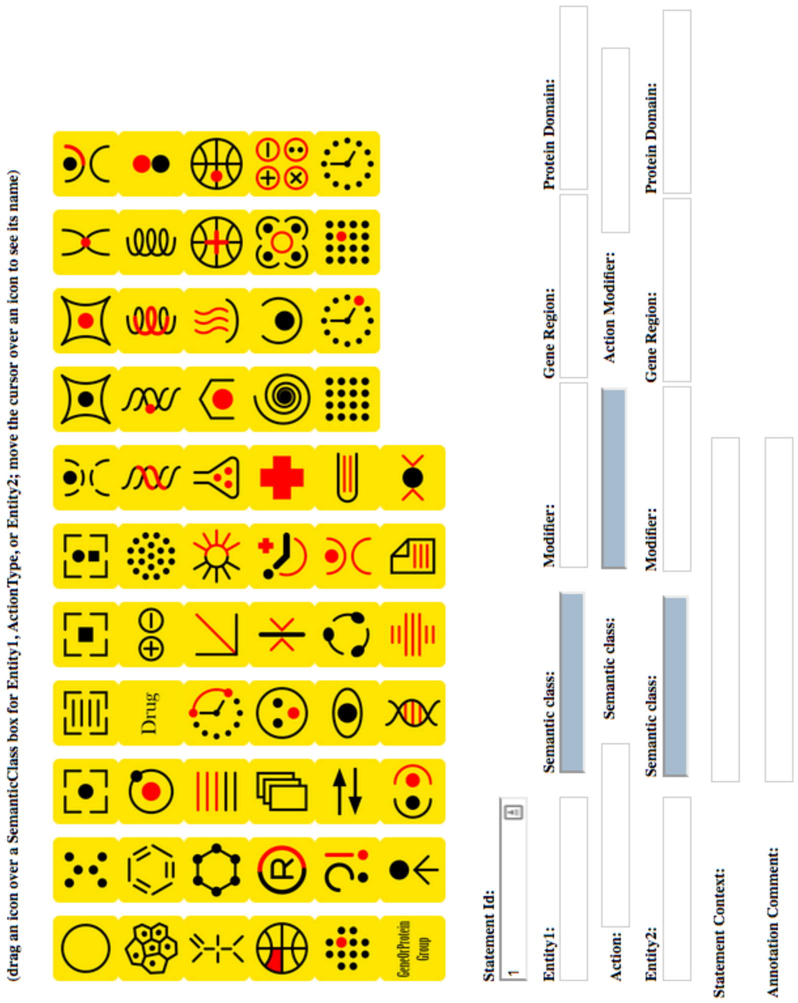


FIGURE 2. **Web annotation tools**

| id | mod1 | g1 | p1 | entity1 | sem1 | action modifier | action | sem action | mod2 | g2 | p2 | entity2 | sem2 | context | user | editor |
|---|---|---|---|---|---|---|---|---|---|---|---|---|---|---|---|---|
| Statement[1] | | | | Sp1 | GP | | bind | | | promoter | | p21WAF1/CIP1 | GP | in high Ca2+-treated NHK cells | res | cle |
| Statement[2] | | | | NFAT1 | GP | | bind | | | promoter | | p21WAF1/CIP1 | GP | in high Ca2+-treated NHK cells | res | cle |
| Statement[3] | | | | CsA | Chemical | | abrogate | | | | | statement 1 | Process | | res | cle |
| Statement[4] | | | | CsA | Chemical | | is | | | | | Cyclosporin A | Chemical | | res | cle |
| Statement[5] | | | | CsA | Chemical | | abrogate | | | | | statement 2 | Process | | res | cle |
| Statement[6] | | | | statement 3 | Process | | restore | | binding of | | | KLF16 | GP | | res | cle |
| Statement[7] | | | | statement 5 | Process | | restore | | binding of | | | KLF16 | GP | | res | cle |
| Statement[1] | | | | Sp1 | GP | | bind | | | promoter | | p21WAF1/CIP1 | GP | in high Ca2+-treated NHK cells | sub | cle |
| Statement[2] | | | | NFAT1 | GP | | bind | | | promoter | | p21WAF1/CIP1 | GP | in high Ca2+-treated NHK cells | sub | cle |
| Statement[3] | | | | CsA | Chemical | | abrogates | | | | | statement 1 | Process | | sub | cle |
| Statement[4] | | | | CsA | Chemical | | abrogates | | | | | statement 2 | Process | | sub | cle |
| Statement[5] | | | | CsA | Chemical | | is | | | | | Cyclosporin A | Chemical | | sub | cle |
| Statement[6] | | | | statement 3 | Process | | restore | | binding of | | | KLF16 | GP | | sub | cle |
| Statement[7] | | | | statement 4 | Process | | restore | | binding of | | | KLF16 | GP | | sub | cle |

FIGURE 3. **Web Annotation Result Example**
