## Supplementary material for "NERO: A Biomedical Named-entity (Recognition) Ontology with a Large, Annotated Corpus Reveals Meaningful Associations Through Text Embedding": Ontology Corpus Annotation Guidelines

### NERO: a biomedical Named Entity (Recognition) Ontology: Annotation Guidelines

[1] *The Committee on Genetics, Genomics, and Systems Biology, University of Chicago, Chicago, IL 60637, US;*
[2] *The Institute of Genomics and Systems Biology, University of Chicago, Chicago, IL 60637, US;*
[3] *School of Computer Science, University of Manchester, M13 9PL, UK;*
[4] *Institute of Medical Bioinformatics, University of Göttingen, Goldschmidtstrasse 1, 37077 Göttingen, Germany.*
[5] *Computational Bioscience Research Center; Computer, Electrical and Mathematical Sciences and Engineering Division; King Abdullah University of Science and Technology (KAUST) Thuwal, 23955, Saudi Arabia;*
[6] *Goldsmiths, University of London, 8 Lewisham Way, New Cross, London SE14 6NW, UK;*
[7] *Department of Chemical Engineering and Biotechnology, University of Cambridge, Philippa Fawcett Dr, Cambridge CB3 0AS, United Kingdom Alan Turing Institute, 96 Euston Rd, Somers Town, London NW1 2DB, United Kingdom Department of Biology and Biological Engineering, Chalmers University of Technology, SE-412 96 Göteborg, Sweden;*
[8] *National Centre for Text Mining, University of Manchester, M1 7DN, UK;*
[9] *The Information Sciences Institute, University of Southern California, Marina del Rey, CA 90089, US;*
[10] *geneXplain GmbH, Am Exer19b, 38302 Wolfenbüttel, Germany*
[11] *Knowledge Lab, Department of Sociology, University of Chicago, IL 60637, US;*
[12] *Department of Medicine and Department of Human Genetics, University of Chicago, Chicago, IL 60637, US;*

#### Contents

**1 Objective** 2

**2 NERO Ontology** 2

**3 Annotation** 3

**4 References** 17

### 1 Objective

We have constructed a new ontology specifically for annotating text entities, trying to minimize unwarranted arbitrary assignments of semantic labels by annotators. Using this ontology, we annotated a large biomedical corpus to enable a broad spectrum of natural language processing and biomedical machine learning tasks. Our corpus differs from previous efforts in several significant aspects. The Named Entity Recognition Ontology (NERO) and our annotated corpus aim to encompass all entity types that might occur in biomedical literature. In addition to Named Entities, the ontology captures *events* representing a spectrum of relationships between biomedical concepts.

### 2 NERO Ontology

The topic area of the Named Entity Recognition Ontology (NERO) is the lexical representation of entities, rather than the entities themselves. For example, we want vocabulary to represent the set of protein names found in a text, rather than the protein that information content represents. Thus, the main aim of NERO is to enable text annotators or text annotation tools to mark up the a text's lexical content as to the nature of that lexical entity.

For example, in the sentence:

> Activation of NF-$\kappa$B2 and RelB was found in 53.7 and 49.2 % of the 121 ER+ tumours analyzed, with similar levels to ER-breast tumours analysed in parallel for comparisons. (1)

Here, NF-$\kappa$B2 and RelB can be either a gene or protein.

In gene and protein naming conventions, italics is used for genes and mRNAs and normal text is used for proteins. More specifically, human genes and proteins are all capitalized. Mice and rat gene symbols have the first letter capitalized, while protein symbols are all upper-case. In contrast, for flies, both gene and protein symbols can begin with an upper-case letter. However, researchers do not follow these naming conventions strictly and often use the same symbol to represent both a gene encoded for a protein and the protein itself.

An annotator, if forced to commit to either a gene or a protein, risks mis-annotating. Enabling an annotator to commit less strongly by annotating these lexical entities as 'gene or protein named entity' avoids such a risk, but still allows annotations to be made and queries posed and answered.

In NERO, we would like to cover all entity types that might occur in biomedical articles. We start, however, with entities around molecules and their interactions within a cell, their link to disease, and the machinery or tests used to investigate these entities.

Thus, the basic competencies for NERO are:

1. Provide vocabulary for annotating the entities covered in the scope outlined above.

2. Provide abstractions of the lexical items such that annotators can commit to an annotation with an appropriate confidence level.

3. To include knowledge about which biological or domain entity a given text entity represents.

NERO is authored in OWL DL using the **protege** 4 authoring environment.NERO may be downloaded with a license. NERO is a simple ontology; it is not axiomatized highly; it only requires a simple taxonomy to fulfill the competencies above. We use a naming convention in which all class labels end with the suffix 'entity.' Labels also capitalize the initial letter.

NERO covers text entities and hence *DomainEntity*–and all semantic ambiguous classes–sits around the NERO's root. The basic division thereafter is into *TextEntity* and *AbstractEntity*, where *TextEntity* further split into *NamedEntity*, *NamedEntityGroup*, *Relationship* and *Pronoun*. The pronouns amount to a set of commonly occurring English pronouns.

After *NamedEntity*, the hierarchy essentially reflects that which may be seen in many descriptions of biological entities, rather than in the lexical representation of those entities. NERO differs in cases such as *'GeneOrProtein'*, which subsumes both *Gene* and *Protein* using the following axiom: *EquivalentTo: 'Gene' or 'Protein'*. There are no biological entities that are either a gene or a protein, but there are lexical entities that are either a gene or a protein. NERO uses this pattern to express ambiguity between various text entities.

Classes in NERO represent information and not the actual biological entities that the information describes. It is, therefore, straight-forward to link between the lexical or informational entity and the biological entity through a relationship such as *'is about'*. So the NERO class *Protein 'is about' some 'protein'* in an ontology such as the Protein Ontology((2)).

### 3 Annotation

**Data Sources and Preperation** The annotation on the corpus was performed by 10 Ph.D.-level annotators with deep experience in biomedical text annotation or biomedical research.Each annotator was first trained on a practise set of 200-300 sentences before moving on to the 'production' annotation stage.The final corpus consists of 35,865 sentences from 8,080 MEDLINE-referenced articles or abstracts. The sentences are selected for annotators randomly.

**Annotation Guidelines** The guidelines for annotation practice have been developed by early annotators and further discussed and finalized. Any changes to the guidelines were discussed thoroughly, and annotators were informed of those changes made in each version.

We aimed to annotate Named Entities relevant to biomedicine as represented in the NERO Ontology. We intended to capture Named Entities at the most specific level on the ontological tree. See Appendix for the complete guidelines.

**Annotation Process and Interface** In order to facilitate the annotation process, we developed a web-based annotation tool. First, annotators read the sentences. Below each sentence is a group of Named Entity classes represented in graphic icons (Figure 1). Annotators then assign a class for each relevant Named Entity by dragging the icon to the Semantic class. To ensure the class consistency, the Semantic class can only be filled using the icon; annotators are not able to enter the Named Entity class manually (it is greyed out). The annotation tool also allows annotators to annotate modifiers for the Named Entities as well. When two Named Entities interact, annotators were able to annotate the action terms.

After the initial annotation, a second annotator may annotate the sentences. Disagreements were discussed (and occasionally resolved) with a third annotator for any remaining discrepancies. To explain this process, we used the following sentence as an example:

> Cyclosporin A (CsA) abrogated the binding of Sp1 and NFAT1 to the p21WAF1/CIP1 promoter in high Ca2+-treated NHK cells, restoring the binding of KLF16, as assayed by a chromatin immunoprecipitation assay. (3)

The annotation results are shown in Figure 2

**Inter-Annotator Agreement** In order to assess the reliability of our annotations, a portion of the corpus was assigned to multiple annotators. We evaluated the process for the following annotation subtasks:

- *Exact* span matches, where two annotators identified exact the same Named Entity text spans.
- *Relaxed* span matches, where Named Entity text spans from two annotators overlap.
- *Exact* concept matches, where within agreed text span, annotators assigned exact same concept class.
- *Parent* concept matches, where the concept class assigned by one annotator is the parent class of the one by the other annotator.
- *Superclass* concept matches, where the two concept classes assigned belong to the same superclass.
- *Ambiguity* concept matches, where one annotator assigned a semantic ambiguous class which includes the concept assigned by the other annotator.

Due to the difficulty in defining the size of negative annotations, instead of $\kappa$ statistic, we reported inter-annotator agreement(IAA) using positive specific agreement or F-measure following the formula from (4).

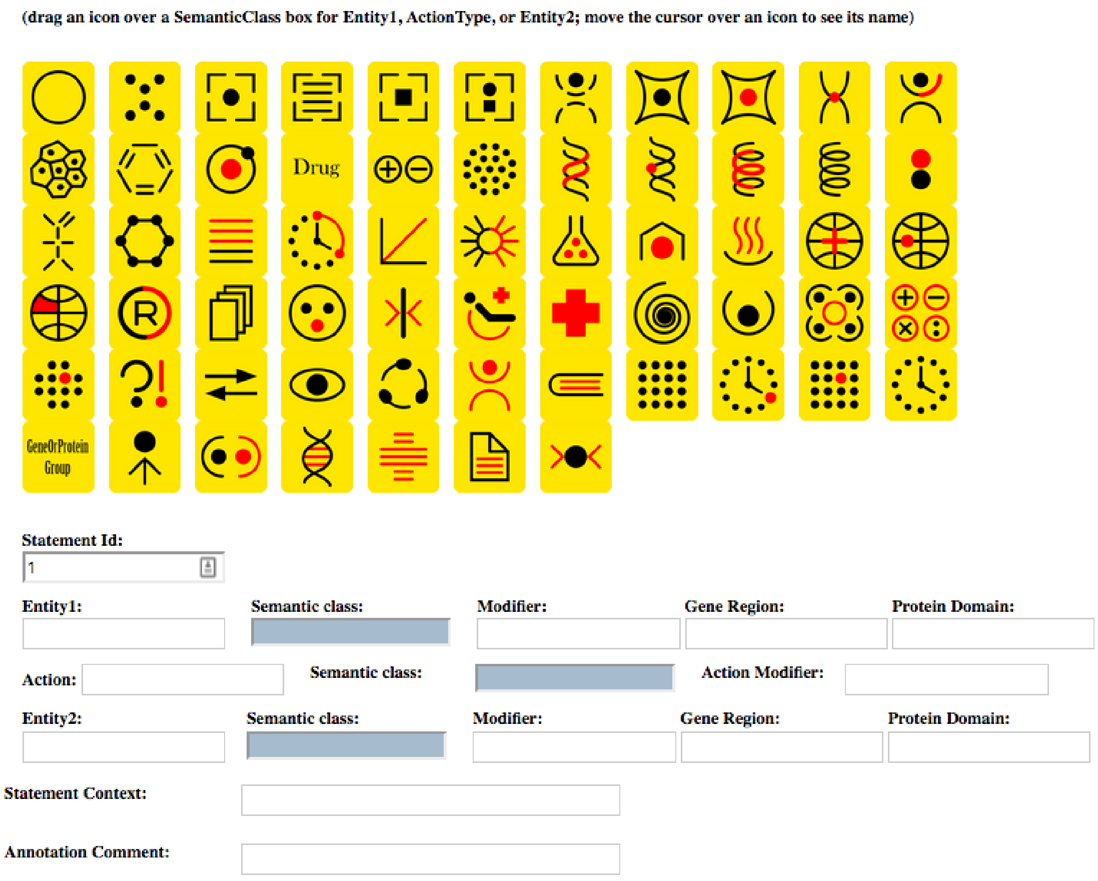

**Figure 1: Web annotation tools**

##### Semantic Classes

***AbstractConcept*** A named entity that can have many meanings. This class is a SUPERCLASS of other classes below. It can be used to define the boundaries of a named entity when more detailed class assignment is difficult. All proper noun phrases that are not better matched as one of the other classes are to be assigned an abstract concept. In those cases where a phrase can be assigned to more than one class, the abstract concept is to be used instead (e.g. Washington could be a person or a location, or Cell could be the name of the Journal or refer to a biological cell – in both cases, *AbstractConcept* should be used.)

| id | mod1 | g1 | p1 | entity1 | sem1 | action modifier | action | sem action | mod2 | g2 | p2 | entity2 | sem2 | context | user | editor |
|---|---|---|---|---|---|---|---|---|---|---|---|---|---|---|---|---|
| Statement[1] | | | | Sp1 | GP | | bind | | | promoter | | p21WAF1/CIP1 | GP | in high Ca2+-treated NHK cells | res | cle |
| Statement[2] | | | | NFAT1 | GP | | bind | | | promoter | | p21WAF1/CIP1 | GP | in high Ca2+-treated NHK cells | res | cle |
| Statement[3] | | | | CsA | Chemical | | abrogate | | | | | statement 1 | Process | | res | cle |
| Statement[4] | | | | CsA | Chemical | | is | | | | | Cyclosporin A | Chemical | | res | cle |
| Statement[5] | | | | CsA | Chemical | | abrogate | | | | | statement 2 | Process | | res | cle |
| Statement[6] | | | | statement 3 | Process | | restore | | binding of | | | KLF16 | GP | | res | cle |
| Statement[7] | | | | statement 5 | Process | | restore | | binding of | | | KLF16 | GP | | res | cle |
| Statement[1] | | | | Sp1 | GP | | bind | | | promoter | | p21WAF1/CIP1 | GP | in high Ca2+-treated NHK cells | sub | cle |
| Statement[2] | | | | NFAT1 | GP | | bind | | | promoter | | p21WAF1/CIP1 | GP | in high Ca2+-treated NHK cells | sub | cle |
| Statement[3] | | | | CsA | Chemical | | abrogates | | | | | statement 1 | Process | | sub | cle |
| Statement[4] | | | | CsA | Chemical | | abrogates | | | | | statement 2 | Process | | sub | cle |
| Statement[5] | | | | CsA | Chemical | | is | | | | | Cyclosporin A | Chemical | | sub | cle |
| Statement[6] | | | | statement 3 | Process | | restore | | binding of | | | KLF16 | GP | | sub | cle |
| Statement[7] | | | | statement 4 | Process | | restore | | binding of | | | KLF16 | GP | | sub | cle |

**Figure 2: Web Annotation Result Example**

***Time*** A period or time point. A calendar time description that includes the year, decade, or century. Other phrases that describe a duration or time related concept are also in this class.

- Polychlorinated biphenyls (PCBs) were measured in the air and water over the Hudson River Estuary during six intensive field campaigns from December 1999 to April 2001.
- The international skeletal society meeting, Budapest 2007: special scientific and radiological focus program, tuesday, 9 october 2007.
- Alois Alzheimer (1864 – 1915) presented the first case of a patient with symptoms of a disease that later would be called Alzheimer's disease.

***GeographicalLocation*** A proper name of a geographical location.

- Both poles of Mars are hidden beneath caps of layered ice.
- Polychlorinated biphenyls (PCBs) were measured in the air and water over the Hudson River Estuary during six intensive field campaigns from December 1999 to April 2001.
- Hospitalizations of patients with acute rheumatic fever were significantly more common in the Northeast and less common in the South.

- **Nested geographic** Thirty-eight patients with chronic heart failure, age 57+/-2 years, New York Heart Association classification II-III, were assigned to either a high intensity training group (n=15, age 53+/-2 years) exercised at 60% of sustained maximal inspiratory pressure, or a low intensity training group (n=23, age 59+/-2 years), exercised at 15% of sustained maximal inspiratory pressure, three times per week for 10 weeks.

***UnproperNamedGeographicalLocation*** A geographical location that is not a proper name.

- Patterns of bacterial diversity across a range of Antarctic terrestrial habitats.
- The net H(+) production associated with Al and Fe transformations was 252 and 1meqm(-2)yr(-1) (on the lake area basis), respectively, reflecting fluxes of ionic, organic, and particulate forms into and out of the lake and the pH gradient between the inlet and outlet.
- Both poles of Mars are hidden beneath caps of layered ice.
- Effect of restricted suckling on milk yield, milk composition and udder health in cows and behaviour and weight gain in calves, in dual-purpose cattle in the tropics.

***PersonGroup*** A proper name of an association of individuals, including companies, clubs, political organizations, government branches, or other entities, as well as groups by character, of people that share certain characteristics such as profession, gender, nationality, or disease.

- Thirty-eight patients with chronic heart failure, age 57+/-2 years, New York Heart Association classification II-III, were assigned to either a high-intensity training group (n=15, age 53+/-2 years) exercised at 60% of sustained maximal inspiratory pressure, or a low-intensity training group (n=23, age 59+/-2 years), exercised at 15% of sustained maximal inspiratory pressure, three times per week for 10 weeks.
- Residents of this valley are predominantly nonsmoking members of the Church of Jesus Christ of Latter-day Saints (Mormons).
- Epigenomics and disease, tenth anniversary winter meeting of the UK Molecular Epidemiology Group (MEG), The Royal Statistical Society, London, UK, 8th December 2006.
- lawyers, physicians, journalists
- diabetics, hurricane victims
- Americans, Russians, Spaniards
- educated people, people of good will

***Person*** A proper name of an individual person.

- Arsenic speciation of two specimens of Napoleon's hair.
- Ten registered Democrats and ten registered Republicans were scanned in an event-related functional MRI paradigm while viewing pictures of the faces of George Bush, John Kerry, and Ralph Nader during the 2004 United States presidential campaign.
- Alois Alzheimer (1864-1915) presented the first case of a patient with symptoms of a disease that later would be called Alzheimer's disease.

***Organism*** An organism, including plant, alga, fungus, virus, bacterium, archaeon, and animal. Including developmental and post-mortem stages. Also covers humans as organisms.

- Arsenic speciation of two specimens of Napoleon's hair.
- Deficiency in recapitulation of stage-specific embryonic gene transcription in two-cell stage cloned mouse embryos.
- Interference competition between introduced black rats and endemic Galápagos rice rats.
- Effect of restricted suckling on milk yield, milk composition, and udder health in cows and behaviour and weight gain in calves, in dual-purpose cattle in the tropics.

Does not cover humans as individual persons (which would fall under Person.)

***AnatomicalPart*** A multi-cellular organization or location of an organ. It includes body parts, body location, body regions, body fluids, organs, organ components, tissue, anatomical structure.

- 
- Ten registered Democrats and ten registered Republicans were scanned in an event-related functional MRI paradigm while viewing pictures of the faces of George Bush, John Kerry, and Ralph Nader during the 2004 United States presidential campaign.
- Two months later, during follow-up, a chest X-ray and computed tomography documented a coin lesion of the upper left lung, confirmed by positron emission tomography.
- Non-suicidal self-injury is the intentional destruction of body tissue without suicidal intent and for purposes not socially sanctioned.

Does not include organisms living within organisms.

***Cell*** A cell or cell line that is not an organism.

- Children with autoimmune disease and CNS injury also exhibited abnormal T-cell responses against multiple cow-milk proteins.
- Neonatal and adult microglia cross-present exogenous antigens.
- The effects of the LABAs salmeterol and formoterol on the synthesis of soluble interleukin-8 (IL-8), granulocyte-macrophage colony-stimulating factor (GM-CSF), and vascular endothelial growth factor (VEGF) in the human airway epithelial cell line A549 was investigated in vitro.

***CellularComponent*** A sub-cellular structure that is neither a gene nor a protein nor a nucleic acid structure.

- Putative, full-unit length begomoviral DNA multimers were digested with Nco I and cloned into the plasmid vector pGEM7Zf+.
- Vinculin links integrin receptors to the actin cytoskeleton by binding to talin.

- Caveolae are extremely stable elements of PECs and can be excluded from their cell membrane only in response to the dramatic cell reconstruction observed in FSGS and LGN.
- Changes in cell morphology and cytoskeletal organization are induced by human mitotic checkpoint gene, Bub1.

***GeneOrProtein*** A gene or protein name, including peptides, but excluding partial sequences. This class also includes secondary structures like alpha sheets and beta coils.

- Children with autoimmune disease and CNS injury also exhibited abnormal T-cell responses against multiple cow-milk proteins.
- The effects of the LABAs salmeterol and formoterol on the synthesis of soluble interleukin-8 (IL-8), granulocyte-macrophage colony-stimulating factor (GM-CSF), and vascular endothelial growth factor (VEGF) in the human airway epithelial cell line A549 was investigated in vitro.
- Liposomes incorporating a Plasmodium amino acid sequence target heparan sulfate binding sites in liver.

***GeneOrProteinGroup*** A group of proteins or gene clusters

- Gene expression of CYP3A4 , ABC-transporters (MDR1 and MRP1-MRP5), and hPXR in three different human colon carcinoma cell lines.
- Genome sequence analysis of Streptomyces ambofaciens ATCC23877 has revealed numerous secondary metabolite biosynthetic gene clusters, including a giant type I modular polyketide synthase (PKS) gene cluster, which is composed of 25 genes (nine of which encode PKSs) and spans almost 150 kb, making it one of the largest polyketide biosynthetic gene clusters described to date.

***AminoAcid*** An amino acid name or sequence of amino acids. This class also includes small peptides that that map to a part of a protein-coding gene or single amino acids.

- Liposomes incorporating a Plasmodium amino acid sequence target heparan sulfate binding sites in liver.
- Cleaving Ala(444)-Ala(445) released mini-plasmin with secondary activity to hydrolyze fibrin.
- Mass spectrometry analysis of the Ebola virus soluble glycoprotein sGP identified a rare post-translation modification, C-mannosylation, which was found on tryptophan (W) 288.

Peptides that are stand-alone would fall under Gene-or-Protein.

***NucleicAcid*** A chemical structure that is based on nucleic acids. It includes nucleoside, nucleotide, RNA, DNA, sites in a sequence, and artificially constructed sequences such as vectors or plasmids.

- Putative, full-unit length begomoviral DNA multimers were digested with Nco I and cloned into the plasmid vector pGEM7Zf+.
- The deletion occurred at the consensus cleavage site (3'-A—TTTT-5') without target site duplication.
- The very long telomeres in Sorex granarius (Soricidae, Eulipothyphla) contain ribosomal DNA.

Does not include chromosomes or genes (the latter would fall under Gene-or-Protein).

***Chromosome*** A chromosome, chromosome region, chromosome part, or chromosome position. It does not include chromosome positions that can be considered measure in units (e.g., 300 bp).

- The very long telomeres in Sorex granarius (Soricidae, Eulipothyphla) contain ribosomal DNA.
- No evidence of linkage between 7q33-36 locus (OTSC2) and otosclerosis in seven British Caucasian pedigrees.
- Failure to confirm allelic and haplotypic association between markers at the chromosome 6p22.3 dystrobrevin-binding protein 1 (DTNBP1) locus and schizophrenia.

***NonNucleicAcidNonProteinChemical*** A chemical structure or a material that is not a gene, a protein, an amino acid, a chromosome, or based on nucleic acids. It includes chemical elements, ions, isotopes, organophosphorus compounds, carbohydrates, lipids, pharmacological substances, and drugs. Drugs are recorded under this category, even when the drug's composition substances are unknown.

- Polychlorinated biphenyls (PCBs) were measured in the air and water over the Hudson River Estuary during six intensive field campaigns from December 1999 to April 2001.
- The net H(+) production associated with Al and Fe transformations was 252 and 1meqm(-2)yr(-1) (on the lake area basis), respectively, reflecting fluxes of ionic, organic, and particulate forms into and out of the lake and the pH gradient between the inlet and outlet.
- Neonatal and adult microglia cross-present exogenous antigens.

Does not include food.

***Food*** Food or drink that is not a simple substance (e.g., salt, water) or an organism name (e.g., wheat, pig, rice).

- The 2005 White House Conference on Aging: a new day for White House conferences on aging and food for the future.
- When comparing highest versus lowest levels of intake in multivariable adjusted models, positive associations were observed for several beef / lamb and individual animal protein items, including beef / lamb as a main dish (OR = 2.2, 95% CI: 1.0-4.5), regular hamburger (OR = 1.7, 95% CI: 1.2-2.4), whole eggs (OR = 1.6, 95% CI: 1.0-2.4), butter (OR = 2.4, 95% CI: 1.6-3.5), and total dairy not including butter (OR = 2.6, 95% CI: 1.8-3.7).
- Digestion rate of legume carbohydrates and glycemic index of legume-based meals.

***EnvironmentalFactor*** Environmental factor

- UV light, radiation ...

***Relationship*** Phrases that express or imply a relationship between objects.

- Mathematical, statistical, or logical relationships: correlation, causation, dependency, equality, progression, significant difference, inverse ...
- Comparisons: similarity, dissimilarity, commonality, increased risk
- Kinships: descendant, ancestor, sibling ...

***Process*** A general, organismal, cellular, or chemical process. This includes processes on the organismal level involving whole tissues or groups of cells such as growth and pathogenesis. It includes processes at the cell level or involving sub-cellular components (e.g., organelles), such as differentiation or apoptosis.

- Caveolae are extremely stable elements of PECs and can be excluded from their cell membrane only in response to the dramatic cell reconstruction observed in FSGS and LGN.
- Thyroid hormone receptor-beta (TRbeta1) impairs cell proliferation by the transcriptional inhibition of cyclins D1, E, and A2.
- The irreversible nature of mitotic entry is due to the activation of mitosis specific kinases such as cdk1/cyclin B.
- Since wee1 keeps cdk1/cyclin B inactive during the S and G(2) phases, its activity must be down-regulated for mitotic progression to occur.

***MolecularProcess*** An activity or event at the chemical or molecular level, including macromolecules like genes or proteins.

- Arsenic speciation of two specimens of Napoleon's hair.
- Deficiency in recapitulation of stage-specific embryonic gene transcription in two-cell stage cloned mouse embryos.
- Liposomes incorporating a Plasmodium amino acid sequence target heparan sulfate binding sites in the liver.
- Digestion rate of legume carbohydrates and glycemic index of legume-based meals.
- Thyroid hormone receptor-beta (TRbeta1) impairs cell proliferation by the transcriptional inhibition of cyclins D1, E, and A2.
- Differential intracellular distribution of DNA complexed with polyethylenimine (PEI) and PEI-polyarginine PTD influences exogenous gene expression within live COS-7 cells.
- The net H(+) production associated with Al and Fe transformations was 252 and 1meqm(-2)yr(-1) (on the lake area basis), respectively, reflecting fluxes of ionic, organic, and particulate forms into and out of the lake and the pH gradient between the inlet and outlet.

***BiologicalProcess*** An interaction at the level of cellular components, cells, organs, organisms, or populations.

- Interactions of immune cells with bacterial cells, cell differentiation, cell death, apoptosis ...
- Hormonal regulation, organ formation and growth, blood pressure regulation, immune response ...
- Digestion, circulation, breathing ...

***MedicalFinding*** Processes that can be considered a specific Medical-finding are to be covered here. An objectively measured sign or symptom (patient-reported problem), or a medical description or observation or finding related to the state of an organism, including sign, symptom, laboratory or test result, syndrome, disease, neoplastic process, mental dysfunction, behavioral dysfunction, or medical finding that is not a measure in units.

- Confirmatory factor analysis of the Epworth Sleepiness Scale (ESS) in patients with obstructive sleep apnea.
- Alois Alzheimer (1864-1915) presented the first case of a patient with symptoms of a disease that later would be called Alzheimer's disease.
- Additionally, the high cholesterol levels found in atherosclerosis could modulate host immunity.

- Thirty-eight patients with chronic heart failure, age 57+/-2 years, New York Heart Association classification II-III, were assigned to either a high-intensity training group (n=15, age 53+/-2 years) exercised at 60% of sustained maximal inspiratory pressure, or a low-intensity training group (n=23, age 59+/-2 years), exercised at 15% of sustained maximal inspiratory pressure, three times per week for 10 weeks.

Does not include cellular, sub-cellular, molecular or chemical processes (e.g., apoptosis, glycemic index).

***MedicalProcedureOrDevice*** A laboratory, therapeutic, diagnostic procedure or method.

- Malignant hyperthermia as a complication of general anesthesia in the clinic of maxillofacial surgery.
- Conventional X-ray exposures in a-p and axial projections and an MRI investigation are considered standard parts of the surgical planning, and a CT examination is also performed when bony defects are present.
- Two months later, during follow-up, a chest X-ray and computed tomography documented a coin lesion of the upper left lung, confirmed by positron emission tomography.

A human-made device, including mechanical, electric, or electronic devices.

- Tomorrow's stethoscope: The hand-held ultrasound device?
- The effect of seat belt use on the cervical electromyogram response to whiplash-type impacts.
- Evaluation of a digitally integrated, accelerometer-based activity monitor for the measurement of activity in cats.
- CT

Does not include buildings or other construction or construction parts (e.g., a room), which fall under the class Facility.

***QuantityOrMeasure*** A numeric value with measuring units, or a phrase expressing a concept of quantity, such as score, dose, rate, size, length, weight, and related terms.

- Thirty-eight patients with chronic heart failure, age 57+/-2 years, New York Heart Association classification II-III, were assigned to either a high-intensity training group (n=15, age 53+/-2 years) exercised at 60% of sustained maximal inspiratory pressure, or a low-intensity training group (n=23, age 59+/-2 years), exercised at 15% of sustained maximal inspiratory pressure, three times per week for 10 weeks.
- High fever, shooting pain, inflammation in the throat, elevated blood sugar.

***Facility*** A construction or part of a construction including buildings, bridges, towers, and other man-made edifices.

- The 2005 White House Conference on Aging: A new day for White House conferences on aging and food for the future.
- Effect of hospital volume on outcome of pancreaticoduodenectomy in Italy.
- The huge garbage dump site near the Hsin-Hai Bridge is likely the source of heavy metal pollution.

Note that whole cities fall under *Geographicallocation* instead.

***Journal*** This refers not to an individual copy of the journal, but to the journal as a regularly published source of information. For an individual copy, or article in such a copy, see Publication.

- Cell, PLoS Biology, Bioinformatics, Time, People.

If the context makes it not clear that the word relates to a journal name, the entity will be classified as abstract concept instead.

***Publication*** A paper, manuscript, video, book, diary, note, message, report, letter, journal, etc.

***Language*** Natural and artificial languages, such as English, Spanish, Hebrew, Turkish, Swahili, Fortran, LISP, C++.

***IntellectualProduct*** A patent, idea, concept, hypothesis. The outcome of a mental process. This is not limited to something that might obtain IP-protection, but may include theories, algorithms, conclusions, and the like.

***MentalProcess*** Memory, emotions, thoughts, learning, cognition. Differs from Intellectual-product in that the focus is on the process of thinking or feeling, not on the result of this process.

***ResearchActivity*** Investigation, measurement, validation, MMPI study, running gel, sequencing. Activities that are executed in the process of conducting research. This includes large-scale operations such as clinical trials, as well as individual lab activities such as sequencing. Used instead of the more general *Process*, if it is clear that the process is a research activity. The more specific *MedicalProcedureOrDevice* is applied if it is clear that the research activity is conducted in a medical context. *MentalProcess* is applied if the activity is a mental process instead.

- Confirmatory factor analysis of the Epworth Sleepiness Scale (ESS) in patients with obstructive sleep apnea.

##### Named Entity detections: Classifier performance

We conducted two initial machine learning experiments. Using NERsuite, we conducted 10-fold cross-validation, dividing corpus into training and test subsets. The classification results are presented in Table 1. The overall performance is moderate, with 54.9% precision, 37.3% recall and 43.4% $F_1$. The best performance class is GeneOrProtein with baseline results of 67.0% precision, 65.3% recall, and 66.2% $F_1$ score.

We then trained an additional set of classifiers on our corpus data for the top 20 classes. We randomly choose 90% of the sentences to be the training set, and the remaining 10% to be the test set. We used this model to tag semantic entities in a fresh set of 141,822 PubMed articles. The performance statistics are shown in Table **??**.

The resulting precision is 51% overall while recall is 42% overall with an overall F1 score of 46%. The Precision is 68% (PersonGroup) on the high end and 23% on the low end. Recall performance varied significantly, with 61%(GeneOrProtein) as the highest and 9% as the lowest. Overall for F1 score, *GeneOrProtein* entities were associated with the best performance of NER engine, 66.38%.

| | **Baseline** | | | **Baseline-Dict Features** | | | **Stacking** | | | **Merging** | | |
|---|---|---|---|---|---|---|---|---|---|---|---|---|
| | **P(%)** | **R(%)** | $F_1$(%) | **P(%)** | **R(%)** | $F_1$(%) | **P(%)** | **R(%)** | $F_1$(%) | **P(%)** | **R(%)** | $F_1$(%) |
| Cell | 62.79 | 56.01 | 59.17 | 62.17 | 55.28 | 58.48 | 62.84 | 56.75 | 59.60 | 60.14 | 53.44 | 56.54 |
| CellComponent | 59.01 | 41.40 | 48.58 | 58.98 | 41.88 | 48.90 | 58.13 | 41.29 | 48.19 | 54.75 | 39.61 | 45.91 |
| GeneOrProtein | 67.00 | 65.35 | 66.16 | 67.05 | 65.81 | 66.42 | 67.02 | 66.04 | 66.52 | 68.33 | 63.52 | 65.83 |
| Organism | 71.72 | 55.14 | 62.32 | 71.35 | 57.00 | 63.33 | 71.03 | 55.70 | 62.40 | 69.73 | 52.58 | 59.92 |
| Disease | 69.72 | 54.75 | 61.28 | 69.21 | 55.11 | 61.29 | 70.23 | 56.93 | 62.83 | 68.63 | 50.72 | 58.26 |
| Drug | 64.13 | 40.40 | 49.43 | 64.88 | 42.95 | 51.59 | 62.19 | 42.51 | 50.35 | 59.60 | 44.18 | 50.64 |
| SmallMolecule | 26.84 | 6.04 | 9.77 | 24.09 | 5.57 | 8.94 | 23.70 | 5.79 | 9.17 | 17.94 | 4.13 | 6.67 |
| BiologicalProcess | 46.03 | 26.64 | 33.71 | 46.08 | 27.23 | 34.19 | 46.19 | 27.24 | 34.23 | 45.71 | 21.07 | 28.81 |
| MolecularProcess | 40.67 | 26.01 | 31.70 | 40.64 | 25.78 | 31.52 | 40.92 | 25.90 | 31.70 | 41.19 | 18.80 | 25.79 |
| Gene | 49.35 | 16.490 | 24.32 | 47.59 | 16.17 | 23.6 | 49.94 | 16.76 | 24.62 | 28.81 | 11.73 | 16.36 |
| Protein | 44.17 | 25.72 | 32.49 | 44.91 | 26.22 | 33.09 | 45.10 | 26.48 | 33.35 | 37.25 | 25.27 | 30.10 |
| BodyPart | 64.62 | 49.02 | 55.72 | 65.05 | 50.30 | 56.70 | 65.23 | 50.13 | 56.67 | 66.75 | 42.86 | 52.18 |
| AminoAcid | 47.53 | 22.37 | 30.20 | 48.88 | 23.24 | 31.29 | 45.15 | 21.29 | 28.72 | 48.10 | 21.84 | 29.75 |
| **overall** | 54.89 | 37.33 | 43.45 | 54.68 | 37.89 | 43.80 | 54.44 | 37.91 | 43.72 | 51.30 | 34.60 | 40.52 |

**Table 1: Experimental results for NER evaluated on 10% of the corpus.**
